# An exploration of ambigrammatic sequences in narnaviruses

**DOI:** 10.1101/761890

**Authors:** Joseph L. DeRisi, Greg Huber, Amy Kistler, Hanna Retallack, Michael Wilkinson, David Yllanes

**Affiliations:** Chan Zuckerberg Biohub, 499 Illinois Street, San Francisco, CA 94158, USA; Department of Biochemistry and Biophysics, University of California, San Francisco, California, USA; School of Mathematics and Statistics, The Open University, Walton Hall, Milton Keynes, MK7 6AA, England

## Abstract

Narnaviruses have been described as positive-sense RNA viruses with a remarkably simple genome of ∼ 3 kb, encoding only a highly conserved RNA-dependent RNA polymerase (RdRp). Many narnaviruses, however, are ‘ambigrammatic’ and harbour an additional uninterrupted open reading frame (ORF) covering almost the entire length of the reverse complement strand. No function has been described for this ORF, yet the absence of stops is conserved across diverse narnaviruses, and in every case the codons in the reverse ORF and the RdRp are aligned. The > 3 kb ORF overlap on opposite strands, unprecedented among RNA viruses, motivates an exploration of the constraints imposed or alleviated by the codon alignment. Here, we show that only when the codon frames are aligned can all stop codons be eliminated from the reverse strand by synonymous single-nucleotide substitutions in the RdRp gene, suggesting a mechanism for *de novo* gene creation within a strongly conserved amino-acid sequence. It will be fascinating to explore what implications this coding strategy has for other aspects of narnavirus biology. Beyond narnaviruses, our rapidly expanding catalogue of viral diversity may yet reveal additional examples of this broadly-extensible principle for ambigrammatic-sequence development.

## Introduction

Narnaviruses (a contraction of *naked RNA viruses*) are RNA viruses with a seemingly simple genome^1^. The only manifestation of narnaviral infections documented to date is the presence of large concentrations of the viral RNA in the cytoplasm of the host cell, often detected as double-stranded RNA. These infections were first observed in cultured yeast^2, 3^. Subsequent metagenomic sequencing revealed narnaviruses in other fungi^4^, oomycetes^5^, mosquitoes^6–9^, other arthropods^10^, algae^11^, trypanosomatids^12–14^ and potentially apicomplexans^15^, although the precise host species is not always clear. The known examples of narnaviruses are approximately 3 kb in size and code for a single protein, an RNA-dependent RNA polymerase (RdRp), with the exception of two putative bipartite narnaviruses^12, 14, 15^.

A remarkable feature of some narnaviruses is the existence of an additional large open reading frame (ORF), defined as a region devoid of stop codons, which spans nearly the full length of the reverse complement sequence of the virus genome. We refer to these examples, with large reverse open reading frame (rORF) features, as *ambigrammatic narnaviruses*, where the adjective is derived from *ambigram*, a set of letters with two, orientation-dependent readings^16^. Such large uninterrupted rORFs are very unlikely to occur by chance, since for a typical nucleotide sequence, there are likely to be codons for which the reverse complement read is a stop codon. This is illustrated in Figure 1A, which shows a typical viral genome spanned by a single ORF and where all five alternative reading frames contain many stop codons. In contrast, Figure 1B shows the genetic sequence of Culex narnavirus 1, where one of the reverse reading frames also has an uninterrupted ORF spanning nearly the whole sequence. All known examples of ambigrammatic narnaviruses have the large reverse ORF in a reading frame where the codons are aligned with those in the forward direction^17^ (i.e., in “frame −0” in the conventions of Figure 2). Figure 1C considers an intriguing intermediate case, which will be discussed further below. The existence of these very long rORFs is a surprising observation which demands an exploration.

**Figure 1.**
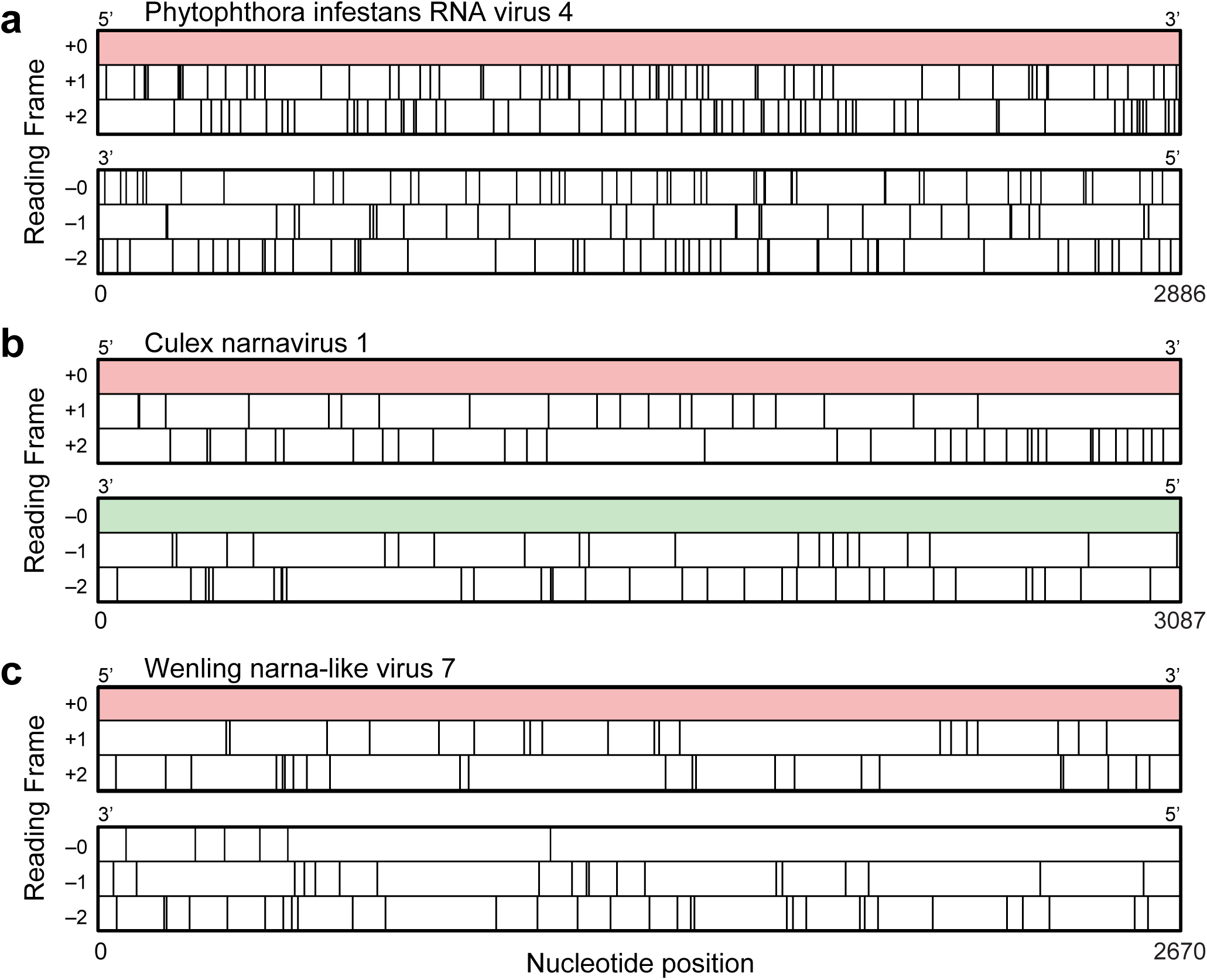
Ambigrammatic sequences in narnaviruses. Coding region for the RNA-dependent RNA polymerase (RdRp) of Phytophthora infestans RNA virus 4 (A), Culex narnavirus 1 (B), and Wenling narna-like virus 7 (C) in the reference +0 frame and all five other reading frames (see Figure 2 for our frame-labelling conventions). Stop codons in each frame are depicted as vertical lines. Large uninterrupted open reading frames (ORFs) are highlighted in colour.

**Figure 2.**
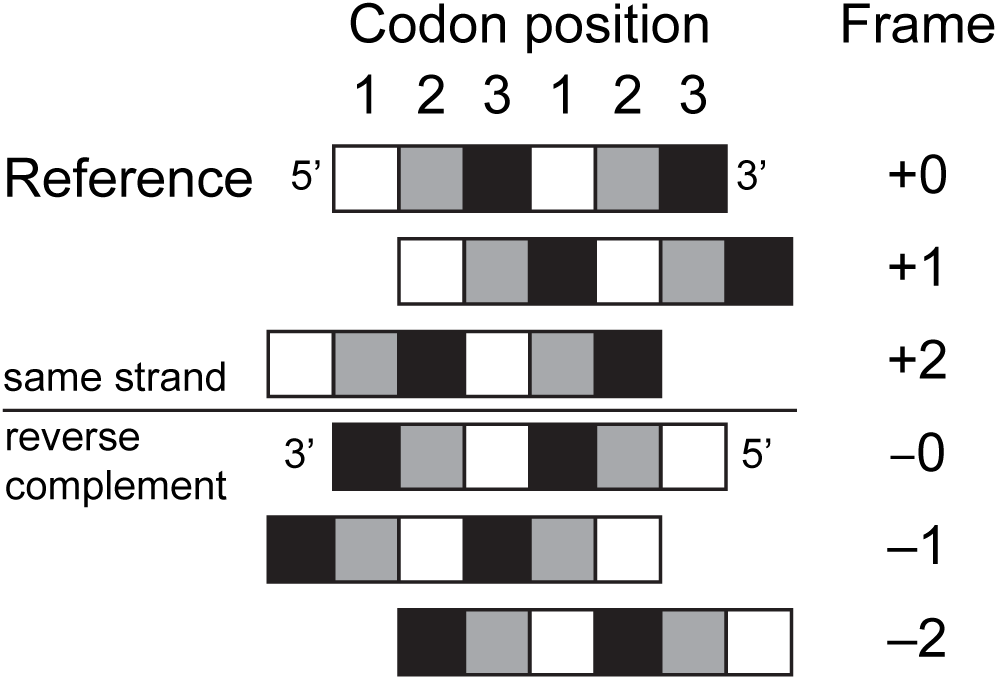
Labelling conventions used in this paper for reading frames.

Interestingly, not all narnaviruses have an ambigrammatic genome (indeed, the example of Figure 1A is also in the *Narnaviridae* family). In agreement with previously reported observations^17^, overlaying the lengths of detectable rORFs on the narnavirus phylogeny shows that ambigrammatic sequences are present in at least two different clades. This finding, illustrated in Figure 3, indicates that the ambigrammatic feature is polyphyletic and may have been gained and lost multiple times in the evolution of this viral family.

**Figure 3.**
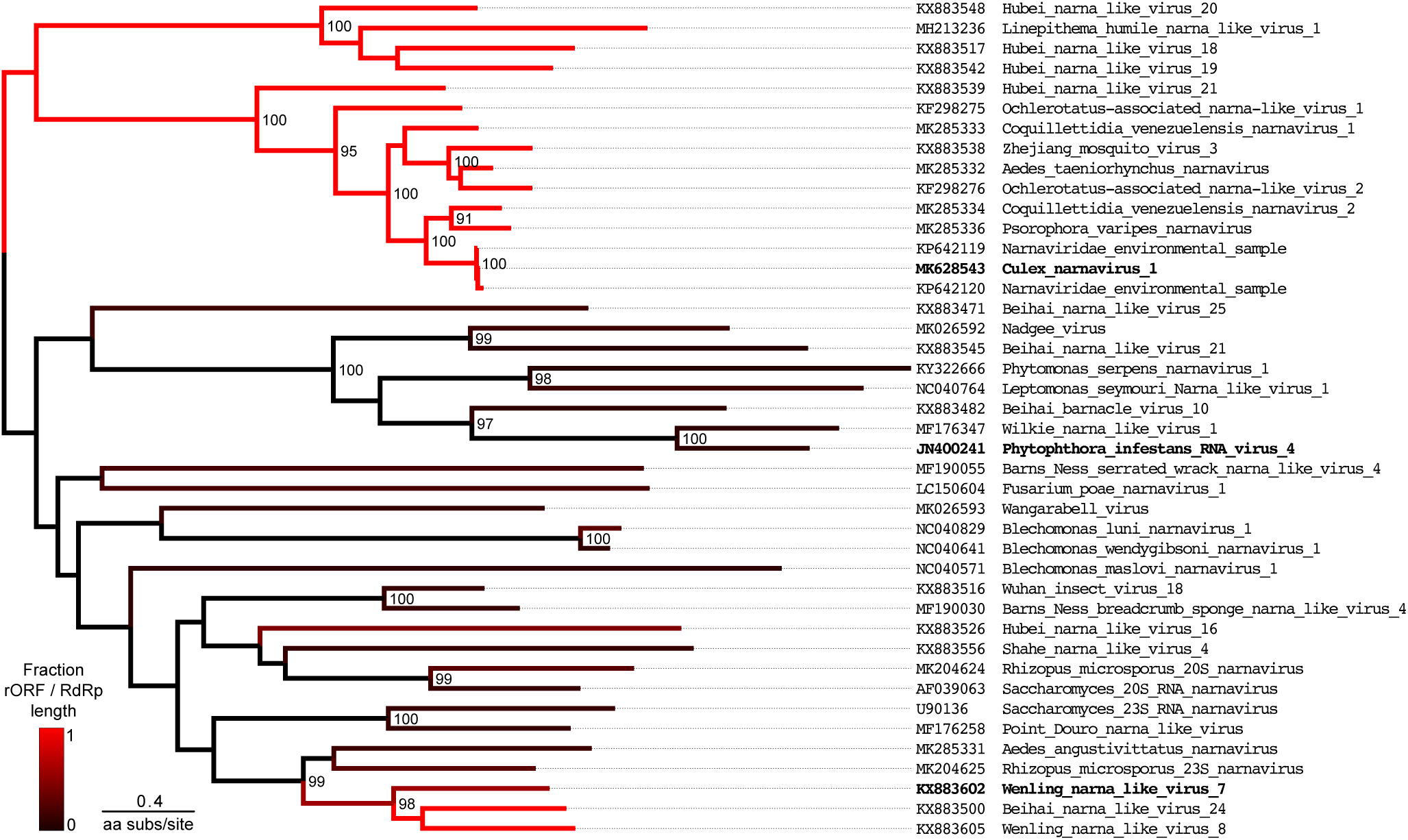
Maximum likelihood tree of amino-acid sequences for RNA-dependent RNA polymerase (RdRp) of 42 representative narnaviruses, identified by homology to the narnaviruses observed in culture, Culex narnavirus 1 and Phytophthora infestans virus 4 (NCBI Blastx^18^). Unrooted tree shown with midpoint rooting for display. Branch colouring indicates the fraction of RdRp coding sequence overlapped by the longest open reading frame (defined as a region uninterrupted by stops) in the reverse complement aligned frame (−0 frame) for sequences at tips (see colour bar, bottom left). The sequence names in bold correspond to those shown in Figure 1. Numbers at nodes indicate bootstrap values (shown when > 80). The branch length is given by the amino-acid substitutions per site, as illustrated by the scale bar.

In this paper we explore how a large open reading frame might arise in the reverse complement sequence of the RNA. While there is an extensive literature on the statistics of codons in overlapping genes (discussed in^19–22^) and on the evolution of overlapping genes in viruses^23–28^, the vast majority of these analyses handle cases of overlapping genes being translated in the same direction, with few papers considering antisense overlaps (see, e.g.,^29^). In contrast, narnaviruses are unique in demonstrating long uninterrupted reading frames in the reverse complement sequence of RNA. Moreover, most analyses of overlapping genes are complicated by the fact that, in the general case, the sequences of both genes can evolve. Analysing cases where one of the overlapping genes is more strongly conserved is very difficult in general, and most earlier works treat the two overlapping genes symmetrically^19^. In this work we adopt a different approach and treat one of the genes as having a fixed amino acid sequence. This could represent a gene with a critical function such as viral polymerases which are often strongly conserved. We suggest that even under these very strong constraints it may be possible for a novel large rORF to arise, but only in *one* of the three possible reverse reading frames. The two alternative forward reading frames, though not directly relevant for experimental observations in narnaviruses, are also discussed for completeness.

We discuss how this mechanism may give rise to a narnaviral genome that is ambigrammatic, in the sense that it can code for two proteins, each translated from one of the two complementary strands of RNA. Finally, we note that ambigrammatic coding in other RNA viruses has not been observed to date, and make some speculative remarks about the potential role of the narnavirus rORF.

## Results

### Synonym Sudoku

Usually, an alternative reading frame for a gene is found to have many stop codons, so that it cannot, therefore, code for a polypeptide or protein of useful length. Let us assume that a nucleotide sequence already codes for a gene which has an essential function. In this case the sequence of amino acids should not be changed. The nucleotide sequence, on the other hand, can still be altered by replacing codons with synonyms (codons which code for the same amino acid). We can ask whether a sequence of single-nucleotide mutations can remove the stop codons that are expected to occur in alternative reading frames, while replacing codons with others that are synonyms in the original reading frame. For some base sequences this will always be possible, but we consider whether *all* of the stop codons in an alternative reading frame can be removed by single-nucleotide mutations in an *arbitrary* polypeptide sequence.

There are five possible alternative reading frames (two in the original strand and three in the reverse complement sequence, see Figure 2). Like a Sudoku puzzle, there is no alternative but to explore all five possibilities in turn. In our discussion, we frame our argument in terms of an RNA genome, so that the base pairings are A=U and G=C, and the stop codons are UAA, UAG, UGA. For the purposes of understanding ambigrammatic sequences observed in narnaviruses, we focus below on the three reverse complement reading frames; the two alternate forward reading frames are discussed in appendix A.

#### Frame −0: aligned complementary reading frames

First, consider the case where the reading frames of the forward and reverse complement sequences have their codons aligned (the −0 frame). In this context, stop codons UAA, UAG, UGA become, respectively, UUA, CUA, UCA in the +0 frame, encoding the amino acids Leu, Leu, Ser. Thus, only instances of leucine and serine in the +0 reading frame can result in stop codons in the −0 reading frame.

We should now consider if synonymous substitution of Leu or Ser codons in the +0 frame can remove the stop codons in the −0 frame. The synonyms of Leu are CU*, UUA, and UUG (where * means any nucleotide). The synonyms of Ser are UC*, AGU, and AGC. Hence, the Leu codon UUA can be transformed to UUG by a single substitution. Similarly, the Leu codon CUA can be transformed to CUU, CUG or CUC by single substitutions, while the Ser codon UCA is transformed to UCU, UCG or UCC by single substitutions. Thus, when frames are aligned, 7 types of single-nucleotide synonymous substitutions in the +0 frame are sufficient to remove stops in the reverse direction.

Ambigrammatic narnavirus sequences have been identified in fungi^17^, where the mitochondrial genetic code uses only two stop codons. The fact that the narnavirus rORF sequences lack all three possible stop codons suggests that translation of the narnavirus rORF in these hosts is not occurring in the mitochondria, unlike viruses of the related Mitovirus genus^1^.

#### Frames −1, −2: staggered complementary reading frames

The cases of staggered reverse complement reading frames are more complex, because a codon in the original +0 reading frame straddles two codons in each of these alternate reading frames. Let us first study the case of the −1 reading frame, where the codons of the reverse reading frame are shifted towards the 3′end (Figure 2). Consider the sequence CUA in the forward direction, with a shift of one base between frames, so we have, using | to denote triplet codon boundaries, *x, y,*… for unspecified bases and 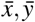,… for their pairing complementary bases:

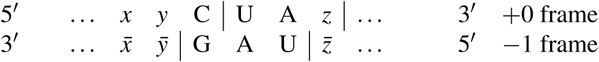

Note that the reversed read UAG is one of the stop codons we wish to avoid, which we could do by changing either UA_*z*_ or *xy*C to a synonym. Let us start with UA_*z*_, which can only be Tyr (either UAU or UAC; the other two values of *y* correspond to stops). UAU and UAC are the only codons that translate to Tyr, so it is not possible to find a synonym of UA_*z*_ that avoids the stop in the reverse sequence. The alternative is to find a synonym of *xy*C. Here we note that if transforming C to U is the only synonymous change, this will still yield a stop codon in the reverse read frame. This occurs if *xy*C represents Asn, Asp, Cys, His, Phe, Tyr, and those Ser codons which begin with AG. Exactly the same restrictions arise from considering the U|UA sequence, and no additional cases of non-removable stops arise from considering U|CA. Thus we find that the following combinations of four +0 codon pairs will prevent an ambigrammatic partner rORF in the −1 frame: (Asn,Tyr), (Asp,Tyr), (Cys, Tyr), (His, Tyr), (Phe, Tyr), (Tyr, Tyr), and some cases of (Ser,Tyr).

In the case of the −2 frame, the codons are shifted to the 5′end. Let us consider when the stop codon in the reverse-read direction is UAA

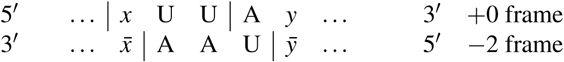

Consider the possible synonym changes to the codons in the +0 frame that will remove the stop codon in the −2 frame. The second codon, beginning with A, can be either Arg, Asn, Ile, Lys, Met, Ser or Thr. Of these, changing the first base can only give a synonym for Arg (AGC and AGT, coding for Ser, yield synonyms by changing two bases). The first codon, ending in UU, can only be Ile, Leu, Phe or Val and it is not possible to obtain a synonym by changing its second base. Finally, it is always possible to obtain a synonym by changing the third base, but, if the first codon is UUU, there is only one synonym, UUC, which also gives a stop in the complementary chain. Considering the case of UC|A yields the same set of excluded combinations, and CU| A does not lead to further examples. We conclude that if the +0 frame contains Phe followed by Asn, Ile, Lys, Met, Ser or Thr; then there is a stop codon in the reverse-read frame which cannot be removed.

#### The potential of synonymous substitutions for generating a new ORF

In conclusion, the −0 reading frame is the only one of the three reverse reading frames where stop codons can always be removed by synonymous single-nucleotide mutations in the +0 original (conserved) amino-acid sequence. This is consistent with the observation that the forward and the reverse open reading frames in ambigrammatic narnaviruses have their codons aligned.

Thus, synonymous substitutions provide a possible route to the generation of a new ORF (as discussed in Appendix A, large ORFs could also be generated in the +2, but not in the +1, reading frame by single-nucleotide synonymous substitutions, although this possibility has not been observed in narnaviruses). We should consider the extent to which this is an effective mechanism for creating new coding potential. In addition to removing stop codons in the −0 reading frame, replacing other codons of the original +0 ORF by synonyms can result in changes to the amino-acid sequence of the rORF, without changing the protein encoded in the +0 frame. It is these synonymous substitutions which allow scope for evolutionary adaptation of the new protein. If the changes in the +0 frame are strictly limited to synonym substitutions, the range of available proteins that can be produced by the new ORFs is quite constrained. We discuss how this can be quantified in the Methods section.

### A hypothesis about the evolution of narnaviruses

First we should consider a null hypothesis, that the rORF (spanning approximately 3 kb) is a chance occurrence. A priori this appears to be extremely unlikely: given that there are three stop codons out of 64 possibilities, we expect that the typical distance between stop codons will be approximately 60 base pairs. If the stop codons are randomly and independently scattered, with mean separation ⟨*N*⟩, the probability of a string of *N* bases containing no stops is expected to be *P*(*N*) = exp(−*N/*⟨*N*⟩). If ⟨*N*⟩ = 60, the probability of finding a sequence of 3 kb lacking a stop codon is approximately exp(−50) ≈ 2 ×10^−22^. The correct value of ⟨*N*⟩ depends upon the distribution of codons, so as a more refined check we examined the distribution of stop codons in each of the five possible alternative frames which arise from choosing a random permutation of the codons of the RdRp gene. Including both ambigrammatic and non-ambigrammatic narnaviruses to generate a null distribution of lengths of ORFs that may overlap the RdRp, we find that the expected probability of a long sequence without a stop codon is indeed well approximated by an exponential distribution, and that the expected probability of an ORF with the observed length is negligible (Figure 4). The scale length ⟨*N*⟩ varies from frame to frame, but is not greatly different from the simple estimate, ⟨*N*⟩ ≈ 60. Of course, even a highly unlikely feature can arise and become fixed in a population. However, there is sufficient variability in the RdRp sequence that stop codons would be expected to arise unless selected against. For instance, the average pairwise identity of the 11 sequences in the clade that includes Culex narnavirus 1 in Figure 3 is only 51 %. These analyses suggests that the chance occurrence and maintenance of a large rORF is highly unlikely, implying that it may offer some evolutionary advantage, broadly speaking.

**Figure 4.**
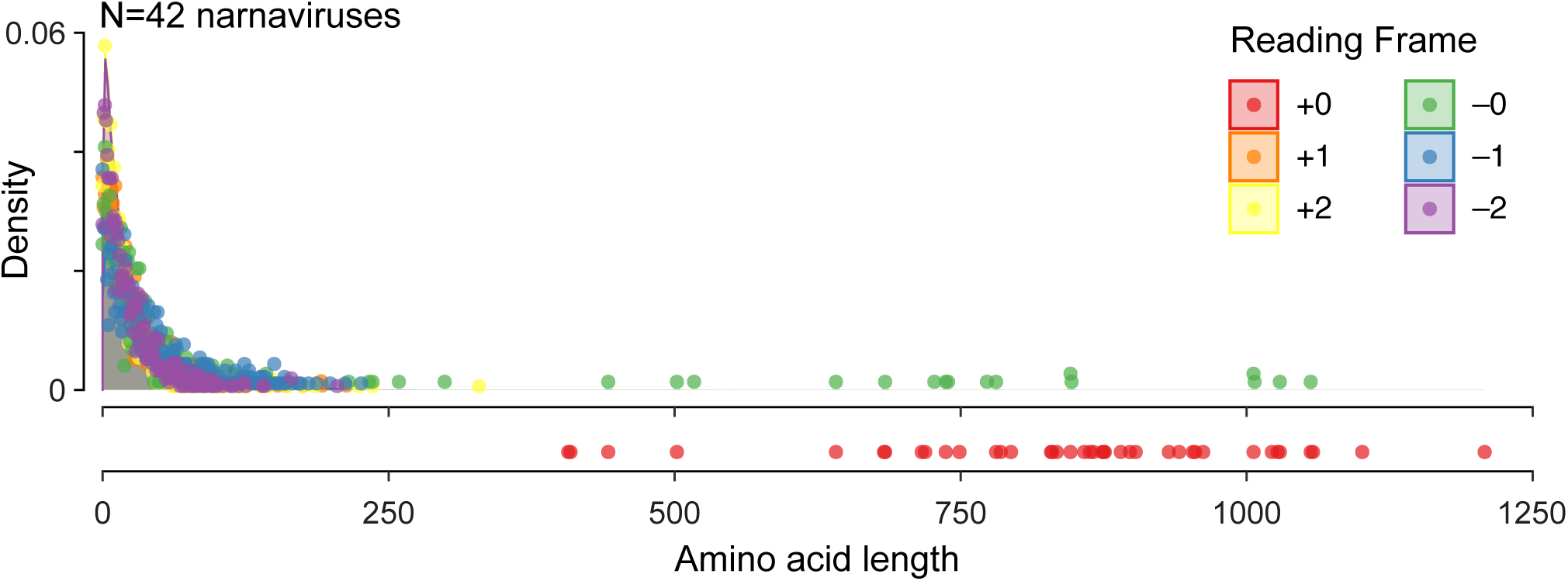
Probability distribution for ORF lengths in narnavirus-like sequences. Shading shows distribution of ORF lengths coloured by reading frame after codon permutation test on RdRp coding sequences of 42 representative narnaviruses as in Figure 3. In brief, codons are randomly re-ordered and then ORF lengths in the 5 alternate frames are calculated (permutation methods as in^30^). Points give lengths of actual ORFs in reference sequences, coloured according to reading frame, with the reference RdRp as +0 frame (red, below). Note that some annotated RdRp coding regions in the database may be fragments of the complete coding sequence.

Given this, it is desirable to speculate how the rORF observed in narnaviruses may have arisen. The virus could have originally existed as a sequence which just coded for the RdRp, with stop codons in the complementary reading frame. It could have evolved by gradually removing the stop codons, and at the same time making other synonym changes in the +0 frame, as the coding sequence lengthens. These mutations could result in progressively longer rORFs, selected by their capacity to increase the fitness of the virus.

The Wenling narna-like virus 7 (Figure 1C), which has a single stop codon in the middle of a large rORF, is intriguing as a potential transitional form in which the rORF is either being gained or lost. If it is being gained, the mechanism in the previous paragraph indicates that the rORF would increase in length incrementally. If the rORF is being lost (for instance, if transfer to exploit a new host removes its advantage), we expect one or more stops randomly scattered in a large ORF. Because this latter picture is more consistent with the Wenling narna-like virus 7 sequence, we speculate that this strain is in the process of losing the large rORF feature. With the presently available data, however, we cannot conclusively support this intuition. Filling out the phylogeny with more sequences could allow us to clarify the potential direction of change.

At this time there is very limited information as to whether the rORF can increase the fitness of the narnavirus. There may be unusual mechanisms in which uninterrupted rORFs alter the translation or protein processing machinery in the cell to the virus’s advantage, even if no protein is made or if the amino acid sequence is not important for its function. For instance, RNA forms of several viruses are thought to be subject to nonsense-mediated decay^31–33^. The narnavirus rORF may be another strategy to increase viral-RNA stability through antagonism of RNA decay pathways or other methods.

Alternatively, if we postulate that the rORF does indeed code for a functional protein, we might speculate on what can be learned from its sequence. A search of the Protein Data Bank (PDB) and NCBI’s translated sequence database (NCBI nr) revealed no sequences with significant homology to the translated rORF. Explorations of secondary structure and other protein features were similarly uninformative, but do not rule out the possibility for a functional protein (see Methods). Perhaps a protein translated from the rORF could enable the narnavirus to evade host-cell defences, allow for movement of the virus between cells, enhance replication by complexing with the genome and RdRp, be required for replication in a particular host species, interact with additional viral elements, or have any of a number of other functions that have not yet been described for this family of viruses.

## Discussion

Motivated by observations of ambigrammatic sequences in narnaviruses, we have shown that an existing ORF can give rise to a large uninterrupted ORF in the reverse complement sequence by synonymous substitutions that preserve the amino-acid sequence of a conserved forward ORF and remove stop codons from the rORF. We find that this mechanism for making ambigrammatic genes only works when the forward and reverse read frames are aligned. These findings are consistent with the observed alignment of overlapping +0 RdRp ORFs and −0 rORFs among many naturally occurring narnavirus sequences.

Any function for the narnavirus rORF remains an intriguing mystery, as does the machinery and processes that may be involved in translating complementary strands of the same RNA sequence. To our knowledge, there are no biologically validated examples of overlapping genes in the reverse complement orientation among RNA-only viruses^26, 27, 34^. While some such overlaps have been predicted^30^, they exhibit neither the overlap length nor the conservation across related strains that is seen among narnaviruses. Indeed, the narnavirus overlap is the longest yet observed among RNA viruses (see, e.g.,^21^). As our observations about the structure of the genetic code are extensible beyond this family, it will be interesting to see whether the explosion in metagenomic sequencing data will reveal more ambigrammatic viruses.

By virtue of packaging, replication, and transmission requirements, viral genomes display a myriad of diverse innovations that provoke us to consider what is possible at the extremes of sequence evolution. Here, the ambigrammatic genes found in some narnaviruses are one such innovation, and their existence likely points to new biology that may be equally as fascinating.

## Methods

### Quantifying the evolutionary space for the companion gene

In a standard gene, a sequence of *N* codons allows for *𝒩* = 20^*N*^ different combinations of amino acids. In the case where a new gene overlaps an existing gene which is perfectly conserved, we must confine our choices to the set of amino acids which correspond to synonyms in the original gene. If the codon for position *j* allows *n*_*j*_ different synonyms, the number of possible amino-acid sequences is

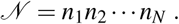

This number grows rapidly with the length of the sequence:

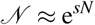

where *s* describes the amount of freedom that the polypeptide sequence has (in physics, it would be termed an *entropy*). For unconstrained evolution we have an entropy per codon equal to

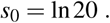

This is a measure of the range of possible proteins that can be constructed in the original gene.

In the cases that we analyse, a new coding sequence in an alternate frame is constrained by the requirement that the amino-acid sequence of the original gene is conserved, so that codons must be chosen from the set of synonyms. The number of possible sequences for the new gene is much smaller, but still grows exponentially with the length of the sequence. Because there are 20 amino acids and 64 codons, there are approximately three possible choices for each codon, so that the entropy of the new gene is (approximately) *s*_1_ ≈ ln 3. Let us consider this more precisely for the case where the new gene is evolving in the −0 frame (that is, reverse complement sequence, with codons aligned). In this case the entropy of the new sequence is

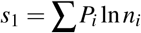

where *P*_*i*_ is the fraction of amino acids of type *i*, and *n*_*i*_ is the number of distinct amino acids which can arise from the reverse complement of each synonym. For example, if the amino acid of the original gene is Ser, there are six possible synonyms (UC*, AGU, AGC). The complementary codons, (UGA, CGA, AGA, GGA, ACU, GCU) represent, respectively Stop, Arg, Arg, Gly, Thr, Ala, so that for *i* = Ser, the number of distinct codons, excluding Stop, is *n*_Ser_ = 4. The corresponding numbers for all of the amino acids are *n*_Phe_ = 2, *n*_Leu_ = 4, *n*_Ile_ = 4, *n*_Val_ = 4, *n*_Ser_ = 4, *n*_Pro_ = 3, *n*_Thr_ = 4, *n*_Ala_ = 4, *n*_Tyr_ = 2, *n*_His_ = 1, *n*_Gln_ = 1, *n*_Asn_ = 2, *n*_Lys_ = 2, *n*_Asp_ = 2, *n*_Glu_ = 2, *n*_Cys_ = 2, *n*_Trp_ = 1, *n*_Arg_ = 4, *n*_Gly_ = 4. Using the codon frequencies *P*_*i*_ determined from the Culex narnavirus 1 sequence, we find *s*_1_ ≈ 1.104, which is remarkably close to the value *s*_1_ ≈ ln 3 ≈ 1.099 estimated in the previous paragraph.

Because *s*_1_ is significantly smaller than *s*_0_, the set of possible sequences which can be coded by the new gene is quite restricted. In particular, in a sequence of length *N* the ratio of the number of possible amino-acid sequences between a standard gene (*𝒩*_0_) and the constrained one (*𝒩*_1_) is

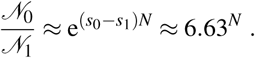

For *N* ∼1000 (as in narnaviruses), this ratio would be ∼ 10^820^. The constraint is eased if we allow changes in amino acids of the original protein as well as synonyms. It is expected that the actual evolution of the virus genome will represent a compromise between conserving the function of the original protein and allowing scope for evolution of the new protein.

### Exploration of the possible secondary structure in the rORF

The translated rORFs of ambigrammatic narnaviruses are predicted to have median *α*-helical and *β*-strand contents of 22 % and 12 % respectively (calculated using JPred4,^35^). This degree of secondary structure is consistent with a structured (or folded) protein, but a significant presence of secondary structure can also be observed in random amino-acid sequences^36^. We further note that the isoelectric point (PI) of the RdRp is high (median 10.4, range 7.7–11.6 for sequences >2 kb in Figure 3), due to a high frequency of Arg, a basic amino acid (median 9.9 %, range 6–13.3 %). Notice that this does not necessarily lead to a high concentration of Arg in the rORF (the codons for Arg are CG*, AGA and AGG, none of which leads to an Arg in the −0 frame). The translated rORFs also have high PIs (median 10.2, range 8.3–11.4), which does not seem to be dictated by the amino-acid composition in the RdRp, and yet is similar to PIs calculated for translations of the −0 frame for non-ambigrammatic narnaviruses (median 10.1, range 6.8–12.7). Basic residues can be involved in binding negatively-charged nucleic acids, but without experimental information we can only speculate on the role for a putative protein translated from the rORF.

## Data availability

All the sequences analysed in this paper are available in online public repositories.

## Acknowledgements

The authors thank Yun S. Song and John Pak for illuminating discussions on yeast genomes and on rORF protein-coding potential, respectively, and Timothy Schlub and Edward Holmes for generously sharing the code used to produce Figure 1. This work and JDR, GH, AK and DY were supported by the Chan Zuckerberg Biohub. HR acknowledges support from the UCSF Medical Scientist Training Program. MW thanks the Chan Zuckerberg Biohub for its hospitality.

## Author contributions statement

MW, DY and GH conceived of the proposal and performed reading frame analysis. HR analysed sequencing data that stimulated this investigation, and generated the figures. JDR and AK provided critical input on viral biology. HR, MW and DY wrote the manuscript with input from all authors.

## A Alternative forward reading frames

In the main text we focused on the possibility of ORFs in the reverse strand, because that was the situation relevant for narnaviruses. However, since the arguments presented in this paper are just based on the genetic code and not specific to these viruses, it is worth considering the alternative reading frames in the forward direction as well.

### A.1 Frame +2

Consider first the case where the new gene is read in the forward direction and left shifted by a single base (frame +2 in the convention of Figure 2). For example, if the codons of the +0 sequence are

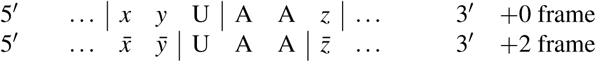

so that there is a stop codon in the new reading frame.

In this case, replacing *xy*U with *xyw, w* ≠ U, removes the stop codon in the new frame. In seven cases (Ala, Arg, Gly, Leu, Pro, Thr, Val) any of the three alternatives for *w* gives a synonym. For another six cases (Asn, Asp, Cys, His, Phe, Tyr), a synonym substitution is also possible but *w* =C is the only option. In the case of Ser, UCU allows three changes but AGU allows only AGC. For Ile there are two possible changes and the remaining amino acids can never have U in the final position. Hence, a synonym for the first codon that removes the stop can always be found.

The same reasoning holds true if AA in the second codon is replaced by AG or GA (the other two possible combinations that would yield a stop in the +2 frame). So, in the case of a left shift, synonym substitution is possible.

### A.2 Frame +1

In the case of a right shift (frame +1), not all stop codons can be removed by a synonym transformation. For example if the codons in the +0 frame are

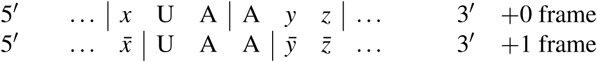

then we want to remove the stop codon UAA by a single-base substitution which gives a synonym, for any choice of *x*. If *x* = U, then we have to find a replacement for the second U or the first A that still codes for Leu. The only possible synonym of Leu that involves changing the second or third letter is UUG, but this is unsatisfactory because UGA is another stop codon. As discussed for the −2 frame, it is only possible to obtain a synonym by changing the A in the second codon if the amino acid is Arg. The same reasoning holds if we consider the UGA stop codon and in the case of the UAG stop codon it is impossible to change the G and obtain a synonym for the second codon in the +0 frame. So the Leu codon UUA followed by any codon beginning with either A or G, except for AGA and AGG (Arg), results in a non-removable stop. With the previous exception, the Leu codon UUG followed by a codon beginning with A results in a stop in the new frame that cannot be removed by a single-nucleotide substitution (although it can be removed by two substitutions, for example UUG → CUG → CUC).

We conclude that it is possible for new ORFs, transcribed in the forward direction, to overlap a perfectly conserved gene, using single-nucleotide mutations to eliminate stops. If we exclude cases requiring more than one substitution, this can only happen in the +2 frame.

